# Revised (β-phenyl) stereochemistry of ultrapotent μ opioid BU72

**DOI:** 10.1101/2020.04.01.020883

**Authors:** Thomas A. Munro

## Abstract

In its crystal structure in complex with the μ opioid receptor (μOR), the opioid BU72 exhibits extreme deviations from the expected geometry.^1^ Three of these involve the phenyl group. There is also unexplained electron density next to the benzylic carbon. Here I show that inverting the benzylic configuration fills this unexplained density and eliminates the phenyl group outliers, along with all but one of the others. I propose that this is the correct structure of BU72.

---

The Protein Data Bank (PDB) validation report for the structure flags twenty outliers in the bond lengths and angles of BU72 (ligand <u>4VO</u>)^2^ with |Z| > 2: that is, more than two standard deviations from the mean observed in crystal structures of subatomic resolution.^3^ In total, a quarter of all evaluated bond lengths and angles are outliers. This gives the structure as a whole root mean square Z (RMSZ) scores of 2.44 for bond lengths and 2.93 for angles. The validation user guide suggests that RMSZ scores > 1 indicate overfitting. In seven cases, |Z| > 5; this is generally considered an extreme deviation,^4^ with a probability of <10^−6^.

The most severe outlier is the near-planar orientation of the phenyl group (Figure 1a). The ideal geometry of the *sp*^*3*^ benzylic carbon is tetrahedral, constrained by a doubly-bridged ring system containing a double bond. Given the extreme implausibility of the modelled conformation (Z = 13.1), the authors speculated that BU72 might have been displaced by a contaminant. This was ruled out by mass spectra of the crystallization mixture, which detected a molecular ion consistent with **1** (or isomers such as **2**), but not the contaminant.

**Figure 1.**
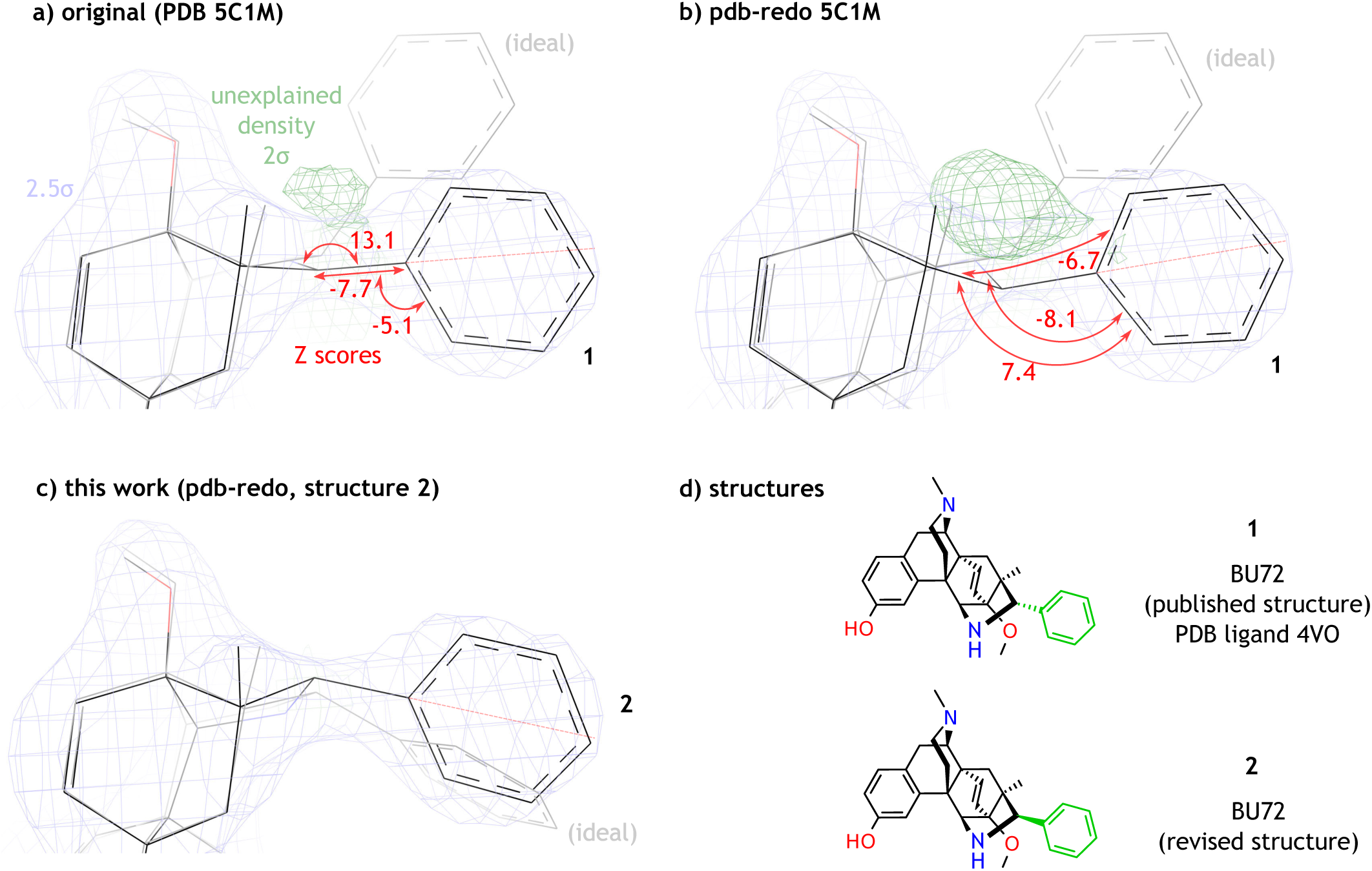
Ligand fit to electron density, phenyl group geometric outliers, and structures 1 and 2. PDB validation Z scores for severe outliers in the phenyl group (in red; torsion Z scores calculated from the XML validation report); ideal structures generated by GRADE server (grey); *2F*o-*F*c map (2.5σ isomesh, blue); *F*o-*F*c omit map (2σ, green).

The phenyl group also exhibits two other extreme outliers. It is attached to the benzylic carbon by an exceptionally short bond, and is bent sharply down from that bond towards the β-face. Although these distortions lower the phenyl group from the ideal conformation into the observed electron density, it remains above center, touching the top but not the bottom of the 2.5σ isomesh (Figure 1a). By contrast, the benzylic carbon lies below center, with a pocket of unexplained density above it.

It is implausible that an ultra-high affinity ligand like BU72 binds in a strained, high-energy conformation, since the energetic penalty of adopting such a conformation reduces affinity.^5^ BU72 has a binding affinity for μOR among the highest known, *K*_i_ = 60 pM^6^ (10 pM with G_i_ protein).^1^ Of 4,979 μOR ligands listed in BindingDB, *K*_i_ < 10 pM for only 9 (0.2%).^7^

The original proposed phenyl configuration of BU72 was based on the nuclear Overhauser effect (nOe): “Irradiation of the 7α-CH_3_ and 5α-H protons enhanced the 2’ and 6’ protons of the phenyl ring, whereas irradiation of the benzylic proton gave no enhancement of … 7α-Me or 5α-H.”^8^ This inference depends on the correct assignment of the signals. However, only a partial listing of ^1^H NMR peaks has been published,^6^ not including the benzylic proton, and the basis for the assignments was not given. Furthermore, the reported nOe enhancement between H-5 and the closest phenyl protons is implausible given their separation in the crystal structure (5.2 and 5.5 Å), beyond the usual detection limit of 4 to 5 Å.^9^ This suggests that some relevant signals may have been misassigned.

The distortion of the phenyl group towards the β-face suggested the β-phenyl epimer **2** as an alternative structure. To test this, the structure of free base **2** and its geometric restraints were generated using GRADE server.^10^ BU72 was then deleted from the model using Coot,^11^ and epimer **2** fitted to the binding site with real-space refinement. The resulting structure was uploaded to PDB_REDO server^12^ for automated refinement, along with the GRADE restraints for **2** and the original reflection data. The resulting model was submitted to the PDB for validation, along with the PDB_REDO re-refinement for **1**. This allows a more direct comparison of the two epimers without differences in modeling and refinement.

In the validation report, epimer **2** shows markedly improved geometry, with only two outliers. The only severe outlier (Z = -7) does not involve the phenyl group. Compared to both models of **1**, the fitted pose for **2** is much closer to the corresponding ideal structure generated by GRADE (Figure 1c, grey). The ideal structure also shows substantial overlap with the observed electron density, unlike **1**. Fit was improved, with no unexplained or excess density around the benzylic carbon, and the phenyl group is better centered in the density. Accordingly, PDB validation metrics are improved (Table 1). I conclude that **2** is the correct structure of BU72.

**Table 1.**
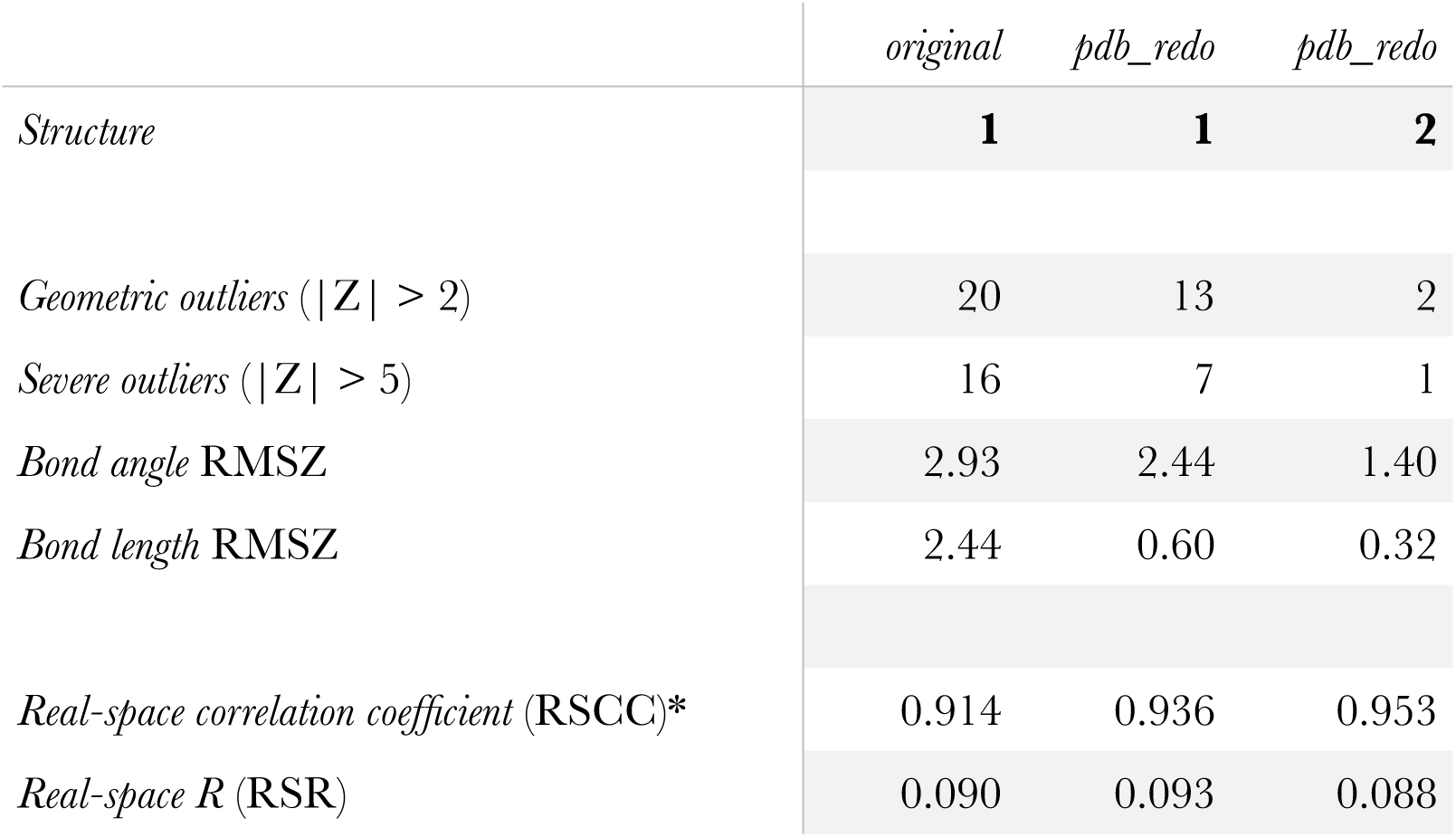
PDB validation metrics for geometry and fit. Lower values are better except for RSCC (*).

## Supporting information

Supporting Information (Coordinates, reflections, ideal structure, validation reports)

## Supporting Information available

Coordinates (mmCif) and reflections (MTZ) for model of **2** with μOR; GRADE ideal structure and restraints for **2**; validation reports for PDB_REDO refinements of **1** and **2** with μOR.

